# Efficient detection of repeating sites to accelerate phylogenetic likelihood calculations

**DOI:** 10.1101/035873

**Authors:** Kassian Kobert, Alexandros Stamatakis, Tomáš Flouri

## Abstract

The phylogenetic likelihood function is the major computational bottleneck in several applications of evolutionary biology such as phylogenetic inference, species delimitation, model selection and divergence times estimation. Given the alignment, a tree and the evolutionary model parameters, the likelihood function computes the conditional likelihood vectors for every node of the tree. Vector entries for which all input data are identical result in redundant likelihood operations which, in turn, yield identical conditional values. Such operations can be omitted for improving run-time and, using appropriate data structures, reducing memory usage. We present a fast, novel method for identifying and omitting such redundant operations in phylogenetic likelihood calculations, and assess the performance improvement and memory saving attained by our method. Using empirical and simulated data sets, we show that a prototype implementation of our method yields up to 10-fold speedups and uses up to 78% less memory than one of the fastest and most highly tuned implementations of the phylogenetic likelihood function currently available. Our method is generic and can seamlessly be integrated into any phylogenetic likelihood implementation.

## 1 Introduction

In phylogenetic analyses, such as maximum likelihood (ML) tree searches or Bayesian inference (BI), the repeated evaluation of the *phylogenetic likelihood function* (PLF) is by far the most costly operation. This is partially due to redundant calculations during the PLF evaluation that can be omitted. Accelerating the PLF is possible by taking into account that (sub-)trees with identical leaf labels (in our case nucleotides), identical branch lengths and the same model parameters always yield the same likelihood score or conditional likelihood values. Therefore, we can save computations by detecting repeating site patterns in the *multiple sequence alignment* (MSA) for a given (sub-)tree topology. From here on, we will refer to those repeating site patterns as *repeats*. Many phylogenetic inference tools such as PhyML [10], RAxML [18], ExaBayes [2], and MrBayes [17] utilize two methods exploiting this property to reduce computations. The first commonly used method consists in evaluating only the likelihood of unique columns of an MSA. Assuming only one set of model parameters for the entire MSA (i.e., unpartitioned analysis), identical sites yield the same likelihood. Therefore, the likelihood can be calculated by assigning a weight to each unique site, which corresponds to the site frequency in the original MSA. In the documentation of PHYLIP [6], Felsenstein refers to this method as *aliasing* (also frequently referred to as *site pattern compression*). The second standard technique for accelerating the PLF at *inner* nodes whose descendants are both *tips* (or *leaves*) is to precompute the conditional likelihood for any combination of two states. Since there is a small, finite number of character states, those pre-computed entries can be stored in a lookup table, and queried when needed, instead of repeatedly recomputing them. These two techniques are standard methods and are incorporated in virtually all PLF implementations providing faster computation times and often, considerable memory savings. In the case of aliasing, memory savings are in the order of 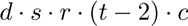, where *d* is the number of duplicate sites, *s* the number of states, *r* the number of rate categories, *t* the number of taxa (tips), and *c* a constant size for storing a conditional likelihood entry (typically 8 bytes for double precision). For example, on a phylogeny of 200 taxa with 100 000 duplicate sites, 4 states (nucleotide data) and 4 rate categories, the memory savings can be as high as 2.5 gigabytes, not to mention the savings in PLF computations.

Apart from the aforementioned standard techniques, there are several studies on improving the run-time of the PLF. Sumner *et al.* presented a method that relies on partial likelihood tensors [20]. There, for each site of the alignment, the nucleotides at each tip node are iteratively included in the calculations. Let *s_i_* be the nucleotide for site *s* at tip node *i*. The values are first calculated for 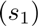, then 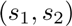, 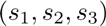 and so on, until 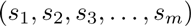 has been processed, where *m* is the number of tip nodes. If the likelihood for another site *s′* with 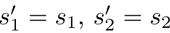 and 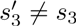 is to be computed, the results for *s* restricted to the first two tip nodes 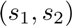 can be reused for this site. A lexicographical sorting of the sites is applied in an attempt to increase the number of operations that can be saved with this method. The authors report run-time improvements for data sets with up to 16 taxa. For more than 16 taxa, the performance of the method is reported to degrade significantly. Additionally, the authors measured the relative speedup of the PLF with respect to their own, unoptimized implementation and not the absolute speedup compared to the fastest implementation available at that time. In [12] the idea of using general site repeats for avoiding redundant PLF operations is mentioned, but dismissed as not practical because of the high book-keeping overhead. Instead, only repeating subtree patterns consisting entirely of gaps are considered since they can be easily identified by using and updating bit vectors, that is, the book-keeping overhead is low. In so-called “gappy” MSAs (alignments with a high percentage of gaps), the authors report a speedup of 25-40% and 65% resp. 68% memory savings on gappy alignments consisting of 81.53% resp. 83.4% gaps (missing data). The underlying data structure used for identifying such repeating subtree sites is called subtree equality vector (SEV) and was originally introduced in [19]. There, only homogeneous subtree columns are considered. That is, a repeat is only stored as such if all nucleotides in that subtree column are identical. This is again done to avoid the perceived complexity associated with finding general (heterogeneous) subtree site repeats. In [19] a speedup of 19-22% is reported for the PLF computation. Similar to [20], the authors of [16] devised a method for accelerating the likelihood computation of a site by storing and reusing the results obtained for a preceding site. Since only the results for one single site (the preceding site) are retained, an appropriate sorting of the sites is required. This column sorting approach is reported to yield speedups in settings where the PLF is evaluated multiple times for the same topology. The authors showed that sorting the sites to maximize the saving potential, can lead to run-time reductions of roughly 10% to over 80%, which corresponds to a more than 5–fold speedup. However, the authors also note, that an ideal algorithm for PLF calculations would reuse all previously computed values from all sites and not just the neighboring ones. Furthermore, the optimal column sorting relies on solving the NP-hard traveling salesman problem and relies on the tree topology. Thus, to construct a polynomial-time algorithm, a search heuristic — that may yield sub-optimal results — is used. This means that, the proposed column sorting may not yield the maximum amount of savings. Another method that focuses on positive selection analysis [21] also deploys a variation of site repeats to accelerate the PLF. The authors implemented their method into a re-designed, optimized version of CodeML (from the PAML package [23]) called FastCodeML and tested its performance against the original CodeML package. Their method, which is specific to codon models and limited to fixed tree topologies, gives speedups of up to 5.8 over the sequential version of CodeML.

Here we show that it is possible to reduce memory requirements and attain a substantial acceleration of the PLF by generalizing the aliasing and tip-tip precomputation techniques. This can be achieved by detecting all conditional likelihood entries at *any* node in the tree, that yield identical likelihood values. Computing these entries only once is sufficient to calculate the overall tree likelihood or any of the omitted (duplicate) entries. The algorithm we present can be applied to both fixed and changing tree topologies. To have a practical application, such an algorithm must exhibit certain properties. First, the overhead incurred by finding repeats must be relatively small such that the overall PLF execution is faster. Second, the book-keeping overhead must be small such that it does not increase the PLF memory footprint. Third, the algorithm and the corresponding data structures must be flexible enough to allow for *partial* tree traversals. When evaluating new tree topologies via some tree rearrangement procedure (e.g., *nearest neighbor joining, subtree pruning and regrafting),* not all conditional likelihood vectors need to be updated. An efficient method for calculating repeats must take this into account and analogously only update the necessary data structures for the partial traversal (i.e., a subset of conditional likelihoods). Thus, the overall goal is to minimize the book-keeping cost for detecting repeats such that the memory usage and run-time is favourable. Furthermore, hardware related issues such as nonlinear cache accesses also need to be considered. For that reason, the absolute speedup of a new algorithm should be determined by using a highly optimized software for PLF calculations and not toy implementations.

We present a new, *simple* algorithm that satisfies the efficiency properties described above; it detects identical sites at *any* node of the phylogenetic tree and not only at the (selected) root, and thus minimizes the number of operations required for likelihood evaluation. It is based on our linear-time and linear-space (on the size of tree) algorithm for detecting repeating patterns in general, unordered, unrooted, n-ary trees [8, 9]. In order to obtain the desired run-time improvements, we present an adapted version of this algorithm for the PLF that reduces book-keeping overhead and relies on two additional properties of phylogenies as opposed to general *multifurcating* (or *n*-ary) trees. First, we assume a *bifurcating* (binary) tree. This assumption can be relaxed to allow multifurcating trees by using a bifurcating tree that arbitrarily resolves the multifurcations. Second, the calculation of the so-called conditional likelihood depends on the transition probability of one state to another. These probabilities are not generally the same for different branches in the tree. Thus, we only consider identical nucleotide patterns to be repeats if they appear at the tips of the same (ordered) subtree. We test the performance of our method against PLL [7] — a library derived from RAxML [18] — which offers one of the most highly optimized PLF implementation. In particular, we show that a prototype implementation of the PLF, that uses our method, consistently outperforms the PLL/RAxML PLF by a factor of 2 to 10. In addition, the memory requirements are significantly lower, with cases where up to 78% **less** memory is required in comparison to RAxML. For the theoretical part of this paper and the sake of simplicity, we assume that genetic sequences only contain the four DNA bases (i.e., `A`, `C`, `G`, `T`). The approach we present can be easily adapted to any number of states (e.g., degenerate DNA characters with gaps or protein sequence data). The data sets we use for benchmarking our method are empirical DNA data sets that *do* contain gaps and ambiguous characters.

## 2 Algorithm

First, we introduce the notation which we will use throughout the paper. A tree 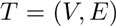 is a connected acyclic graph, where *V* is the set of nodes and *E* the set of *edges* (or *branches*), such that 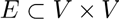. We use the notation 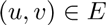 to refer to an edge with end-points 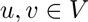 and 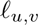 to denote the associated branch length. The set *L(T)* comprises the tip nodes. We use *T_u_* to denote a subtree of a (rooted) tree *T* rooted at node *u*.

### 2.1 The phylogenetic likelihood function

Before we introduce our method, it is necessary to give a brief description of PLF computations. The likelihood is a function of the states σ, the transition probabilities *P* for all branches, and the equilibrium frequencies of the states 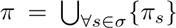. The PLF can be further extended with additional parameters such as variable rates (denoted *C*) of substitution across sites (see for instance [22]). In his seminal paper [4], Felsenstein introduced the *pruning* algorithm for computing the PLF, which is a dynamic programming approach for computing the likelihood of a given tree *T*. The method iteratively computes all *conditional* (or *partial*) likelihoods 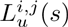, that is, the likelihood for a subtree rooted at node *u*, for site *i* and rate *j*, assuming the state at node *u* is *s*, via a post-order (bottom-up) traversal of the tree. Such a traversal always performs the first computation at a tip node and conducts computations at a node only after both its children were visited. Now, let us assume an MSA of *n* sites (columns) and *m* sequences constructed from *s* states (e.g., 4 for nucleotide data). The conditional likelihood 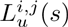 of any tip node 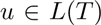 with the sequence 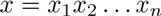 is defined as

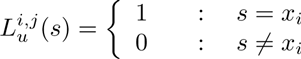

In the case of data with ambiguities or gaps, we replace the conditions 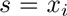 resp. 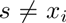 with 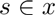 resp. 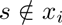. In the case of *inner* nodes 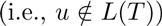, the conditional likelihood for site *i* and rate *j* is defined as

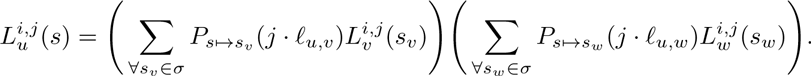

where 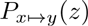 is the probability of state *x* changing to *y* in *z* units of time, and *v* and *w* are the two descendants of *u*. We write the conditional likelihood vector (CLV) entries for all rates and all possible states at a particular site *i* of node *u* as

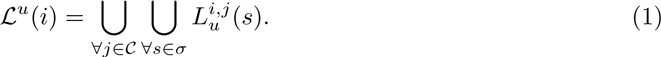

Finally, the overall likelihood *L* for a rooted tree *T* with root node *r* is computed as

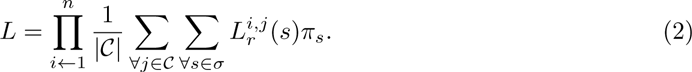

We evaluate the overall likelihood of an unrooted binary tree at branch 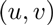 as

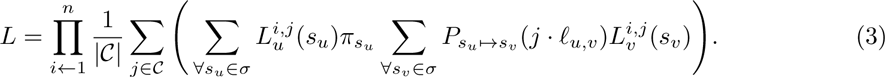

For more details on the PLF, see [5].

### 2.2 Site repeats

Let *T_u_* be the subtree of *T* that presents the evolutionary relationships among the taxa at tip nodes 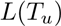. We denote the sequence of the *i*-th taxon 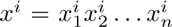. Two sites *j* and *k* are called *repeats* of one another 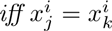 for all taxa *i*, 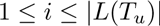, in *T_u_*. Next, we make two observations.

**Observation 1** *If two sites j and k are not repeats in some tree T_u_, then they are not repeats in any tree containing subtree T_u_*.

**Observation 2** *Let u be a node whose two direct descendants (children) are nodes v and w. If two sites j and k are repeats in both T_v_ and T_w_, then j and k are also repeats in T_u_*.

Based on these two observations we can formulate the algorithm for detecting site repeats in binary phylogenetic trees. However, before we formalize the algorithm, let us consider Figure 1 again. From Observations 1 and 2, we see that the only repeating sites at the root node (node *u*), are sites 2 and 5. This is obviously correct, since both have the nucleotide pattern `ACCT` at the tips.

**Figure 1.**
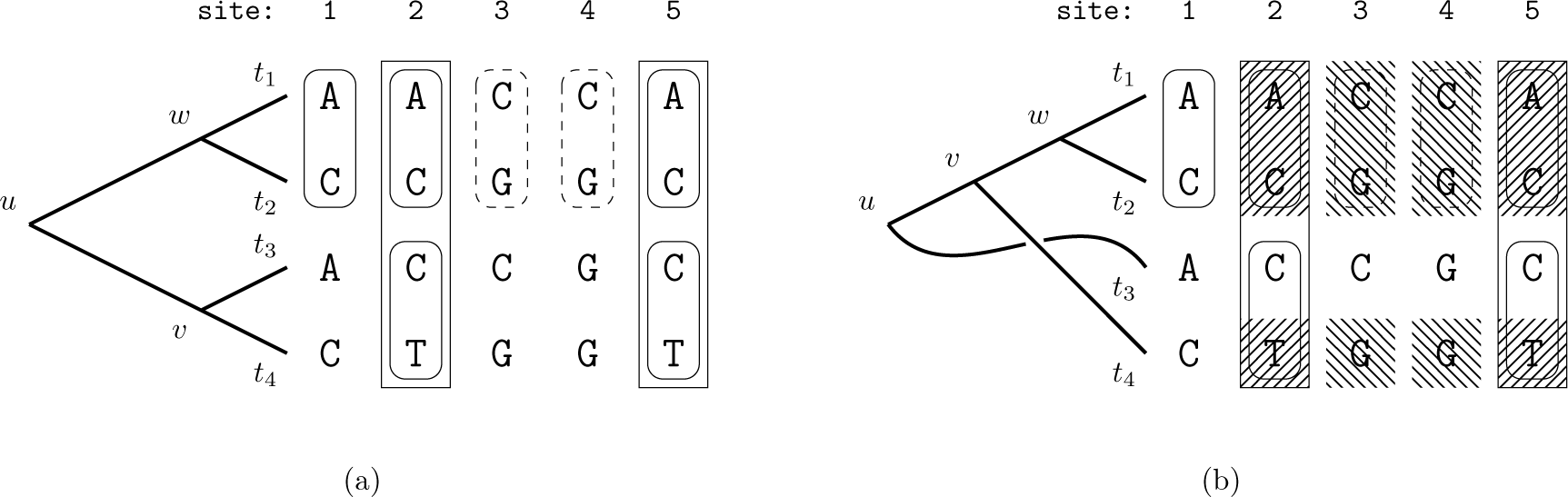
(a) Sites 1,2 and 5 form repeats at node *w* as they share the same pattern `AC`. Another repeating pattern is located at sites 3 and 4 (`CG`) for the same node. Note that, node *u* also induces a subtree with pattern `AC` at the tips. However, since branch lengths can be different than for the subtree rooted at node *w*, the conditional likelihoods may differ as well. Analogously, sites 2 and 5 are site repeats for node *v* as they have the same pattern `CT`, and hence the conditional likelihood is the same for those two sites. Finally, sites 2 and 5 form repeats for node *u* (`ACCT`). (b) Repeats are not necessarily substrings of MSA sites. For this particular tree topology, node *v* has two sets of repeats: sites 2 and 5 (`ACT`) and sites 3 and 4 (`CGG`). The repeats are not contiguous in the alignment columns.

### 2.3 Calculating Repeats

The method we propose identifies site repeats at each node via a bottom-up (post-order) traversal of the tree, meaning that a node is processed once the repeats for its two children have been determined. Tip nodes maintain only the trivial repeats of all sites that show a common character (for DNA, `A`, `C`, `G`, or `T`, respectively). Therefore, the method always starts at an inner node whose two children are tip nodes. By construction, such a node always exists in any binary tree and assuming four nucleotide states, there are 16 possible combinations of homologous nucleotide pairs in the sequences of its two child nodes. To assign a unique identifier to each nucleotide pair, we use a bijective mapping 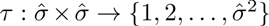, where 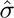 corresponds to the set of *observed* states (4 for nucleotides, or 16 when considering ambiguities and gaps). The problem of identifying repeats is thus reduced to filling, and querying the corresponding entries in a list of CLVs. We outline the identification of repeats for the three possible cases.

**Tip–Tip case**. Assuming that 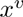 resp. 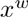 are the sequences at the two children *v* and *w* of the parent node *u*, site *i* of *u* is assigned the *identifier* 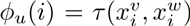. This function assigns the same identifier to sites which are repeats in *T_u_*. Figure 2 illustrates the assignment of identifiers to combinations of nucleotides at the tips for the example in Figure 1. The CLV entries are computed only once for each identifier at the parent node *u*, for example, the first time it is assigned to a site. By Observation 2, if a site *i* is a repeat of site *j* (i.e., both sites are assigned the same identifier), then the method can either (a) copy the CLV from site *j* (run-time saving), or (b) completely omit the likelihood value, since it can always retrieve it from site *j* (run-time *and* memory saving). Furthermore, by Observation 1, we know that each repeat is identified by this method.

**Figure 2.**
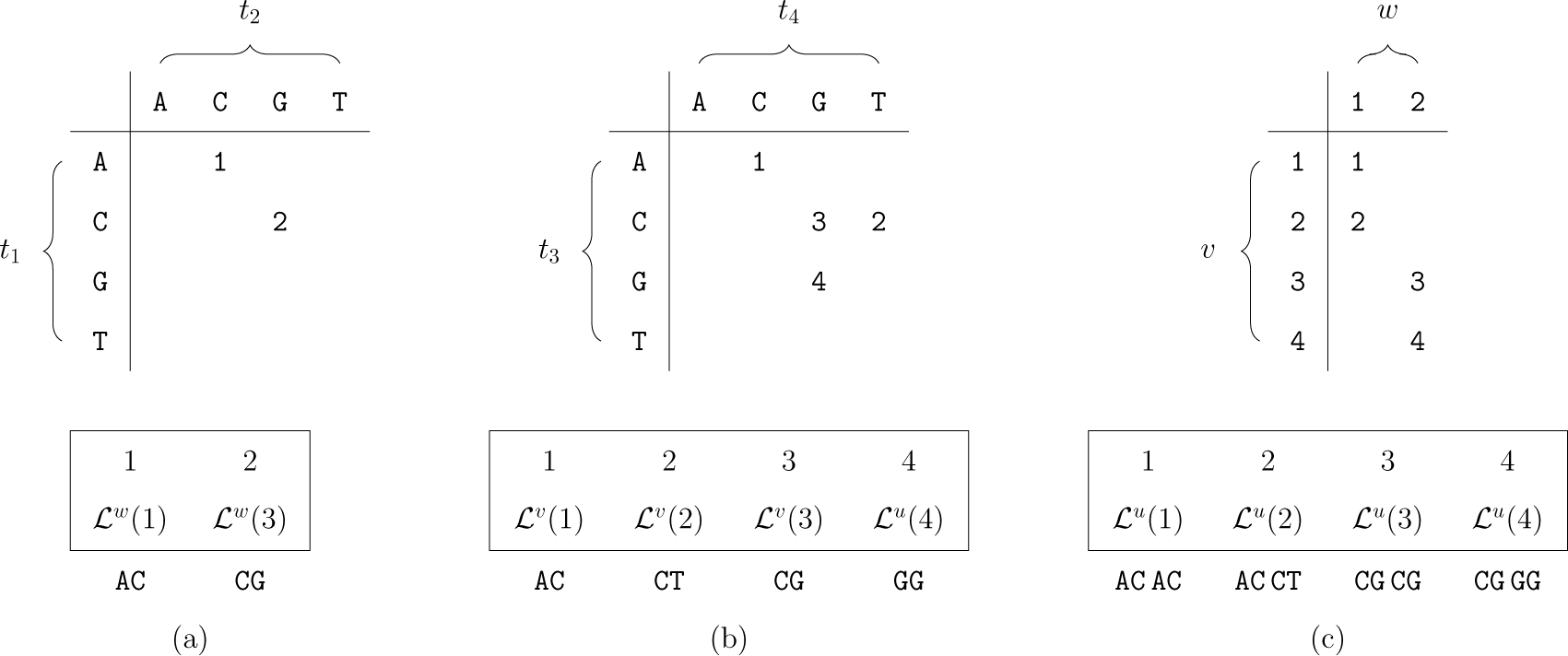
Identifier associations of nodes 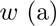, 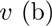, and 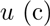 for the tree from Figure 1a. The respective lists at the bottom store the corresponding CLVs that are computed for each unique identifier. Table (a) shows that node w requires two likelihood computations (sites 1 and 3), while the remaining sites are repeats of those two. Tables (b) and (c) show the corresponding information for nodes *v* and *u*.

**Tip–Inner and Inner–Inner cases**. We proceed analogously to detect repeats at nodes for which at least one child node is not a tip. Again, let *u* be the parent node and *v* and *w* the two child nodes for which we already computed all repeats. Further, let 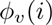 and 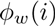 be the respective identifiers of *v* and *w* at site *i*. We define the maximum values of 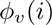 and 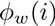 over all sites as *vmax* and *wmax* respectively. Those values represent the number of unique repeats at nodes *v* and *w*. Finding repeats at *u* is again simply a matter of filling the appropriate lists/matrices. Given *vmax* and *wmax*, there are at most *vmax* × *wmax* combinations at the sites. Figure 2c demonstrates the identifier calculation at node *u* for the example tree and MSA from Figure 1. Figure 3 shows the combined overall result.

**Figure 3.**
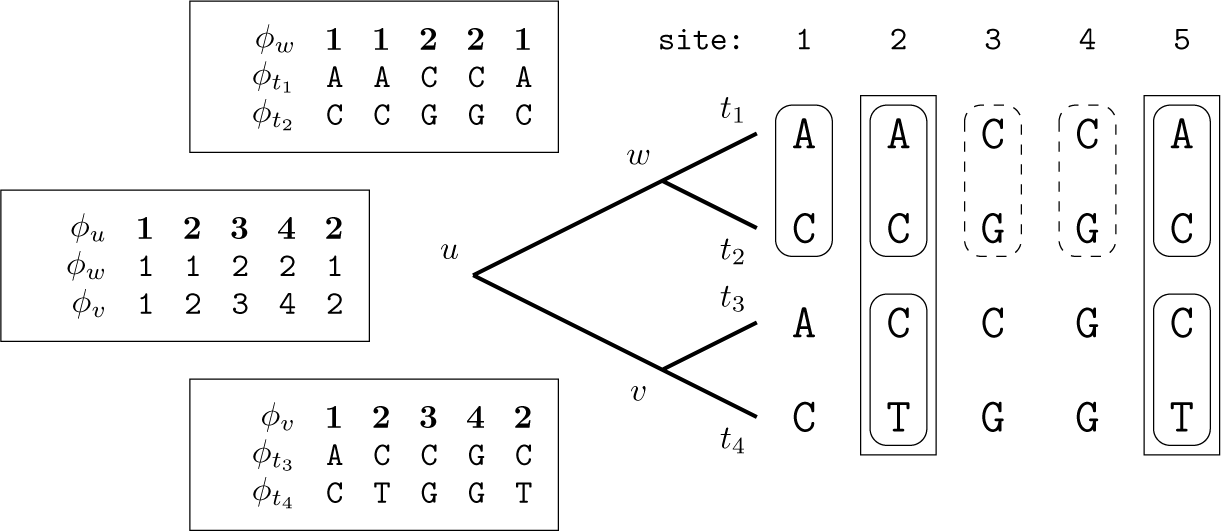
Identifiers (here 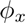) are shown for every site of the alignment at every node in the tree. As we have already observed, sites 2 and 5 are repeats at node *u*, and thus, have been assigned the same identifier. For simplicity, identifiers at tip nodes are represented as nucleotide bases.

Figure 4 outlines algorithm 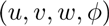, which calculates the CLV for node *u* given child nodes *v* and *w*, by taking into account site repeat information. Using algorithm REPEATS, we can design the complete method which performs a post-order traversal over all nodes of tree *T*. For this, the tip nodes *t_j_* can be assigned constant identifier sequences that correspond to their respective DNA sequences. The actual nucleotides `A`, `C`, `G`, and `T` can simply be mapped to integers. Note that, in most (if not all) phylogenetic inference tools, nucleotides are encoded using the *one-hot* (also called *1 out of N*) encoding, which ensures that the binary representations of their identifiers have exactly one bit set (e.g., 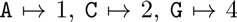 and 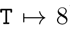). This is beneficial because the identifiers of ambiguities which are typically represented as disjoint unions of nucleotide codes, can be encoded as the bit-wise OR of the identifiers of the respective nucleotides. To simplify the method description, we discard ambiguities and only consider the four nucleotide bases. Hence, we use the encoding

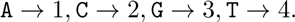

**Figure 4.**
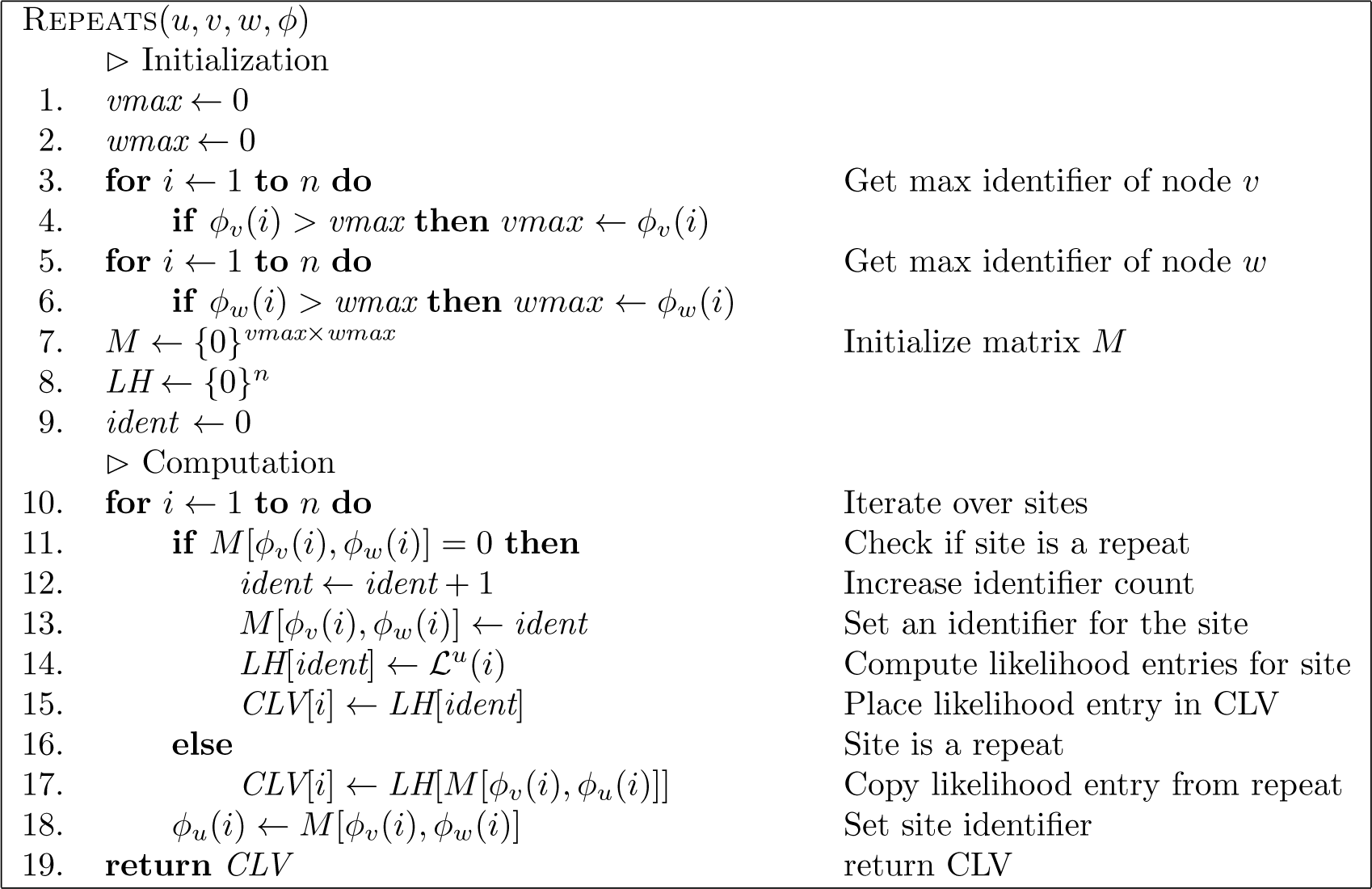
The algorithm to compute the CLV of a parent node *u*. The most costly operation is the calculation of the CLVs, expressed here as 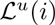 (see Equation 1). The algorithm minimizes the number of calls to this function by taking into account site repeats.

**Lookup Table** Since our focus is on an efficient implementation of the algorithm, we need to consider some technical issues in more detail. First, matrix *M* (defined in algorithm REPEATS) can, in the worst case, become quadratic in size with respect to the number of sites in the alignment. This is unfortunate, since filling *M* affects overall asymptotic run-time. However, in terms of practical space requirements, *M* can be allocated only once and subsequently be reused for each inner node. For that, a linear list *clean* with one entry per MSA site, can be used to keep track of which entries are *valid*, that is, contain identifiers assigned to sites of the current node, and which entries are *invalid* and contain identifiers assigned to the sites of a preceding node. After assigning an identifier *i* to a site of node *u*, which we store in the array *M*, for example at position *d*, we also store the pair (*d*, *u*) in array *clean* at position *i*. Later on, when we process a different node, say *v*, and by chance, decide to assign the same identifier *i* to some site, and again, by chance, the location for which we have to query matrix *M* is *d*, the element *clean[i]* helps us distinguish between valid and invalid records in *M*. Invalid records are equivalent to empty records and are overwritten. Further, in the actual implementation we limit the size of *M* to a constant maximum size. We implement this limit to adapt the impact of the quadratic complexity for filling *M*. In Section 3, Table 2 gives an overview of the size of *M* for different data sets. Since dynamically tuning the size of *M* to the data set can have a negative impact on the run-time and memory performance, the size of *M* is an input parameter. In addition, as *M* grows larger (i.e., we move closer to the root of the tree), it is less likely to encounter repeats in the alignment. Note that, at the CLV of the root, there will be no repeats at all, since they have already been removed by compressing MSA sites into patterns during MSA pre-processing. One may also consider the following alternative view. If *M* is an *n* × *n* matrix, where *n* is the number of sites in the alignment, there can be no repeats, as every site must, by construction, have a unique identifier. If at least two sites were repeats of another, the maximal identifier would be strictly less that *n* and thus, *M* would not be a *n* × *n* matrix. Thus, if the product of maximum identifiers for two child nodes at some node *u* (that is, *vmax* × *wmax*) exceeds our threshold for the size of *M*, we do not calculate repeats any more. Instead, the CLV entries are calculated separately for all sites as in standard PLF implementations. In other words, if calculating repeats becomes disadvantageous, repeat calculations are omitted. This allows to trade repeat detection overhead, against PLF efficiency.

**Memory savings** Notice that, given algorithm REPEATS, not all entries in the CLVs of the child nodes *v* and *w* are needed to calculate the CLV at the parent node *u*. In particular, the CLV entries at site *i* for nodes *v and w* are only needed if the CLV at site *i* must be computed for *u* (see Figure 5). In fact, only the CLVs in array *LH* of algorithm REPEATS must be stored. Let 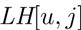 be the CLV computed by Algorithm 4 for node *u* and the site with identifier *j*. Then, the CLV for any site *i* is simply 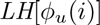. In practice, this observation allows us to reduce the memory footprint of the PLF. Each CLV entry stores more than one single or double precision floating point value. For example, RAxML stores one double precision floating point number per DNA character and per evolutionary rate for each CLV entry. Typically, the Γ model of rate heterogeneity is used (see [22]) with 4 discrete rate categories. Thus, the memory footprint of a standard PLF algorithm for a MSA with *n* sequences of length *m* is 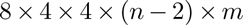 bytes. On the other hand, storing the site identifiers at each node only requires a single, unsigned integer per site. Thus, the memory required for storing CLVs without compression is 4 • 4 = 16 times higher than that of the site identifier list.

**Figure 5.**
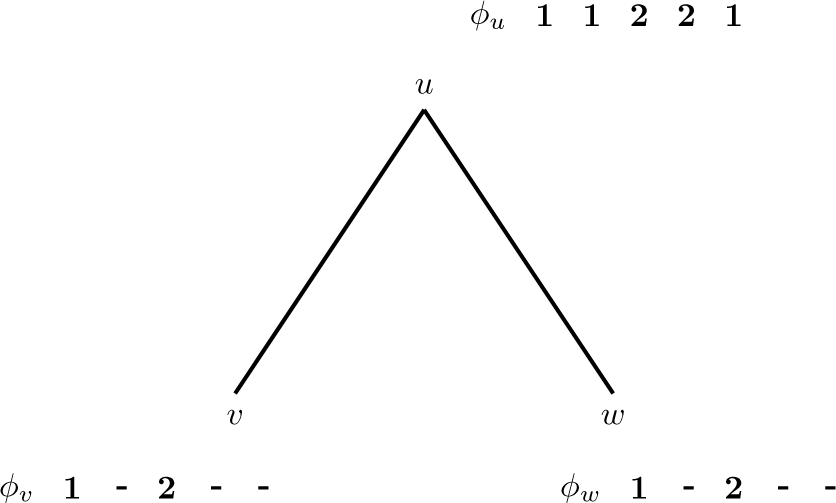
Not all sites are needed for the likelihood calculation at parent node *u*. According to the identifiers of this example, sites 2 and 5 are repeats of site 1, and site 4 is a repeat of site 3. Therefore, the CLVs at sites 2, 4, and 5 do not need to be computed nor stored, as the CLV for sites 2 and 5, and site 4, of node *u* can be copied from sites 1, and 3, respectively.

Thus, despite the fact that we need additional data structures, and hence space for keeping track of the site identifiers at nodes, the memory requirements (if we do not store unnecessary CLV entries) are smaller than those of standard production level tools [7, 18]. While the identifiers are not the only additional data structures required for the actual implementation of the algorithm, the above argument illustrates that storing fewer CLV entries can help to save substantial amounts of RAM. The overall algorithm, with memory savings and a bounded *M*, is given by algorithm REPEATS-FULL in Figure 6. One main difference to the snippet of Figure 4, is the introduction of a new array (*maxid*) which stores the maximal identifier assigned to each of the 2*m* – 1 nodes of the rooted tree *T* (assuming *T* has *m* tip nodes). Thereby, we eliminate the run time 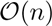 required for finding the maximal identifiers of the two child nodes (lines 3-6 in Figure 4) at the cost of 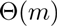 memory. The second difference is that, we can no longer use the original set 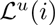 for the CLV entries of a site *i* at a node *u*. This is due to the memory saving technique which omits the computation and storage of unnecessary CLVs as illustrated in Figure 5. The problem is that the CLV of the two children may not reside at entries *i* because repeats might have occurred. Therefore, the new set 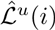 is defined as

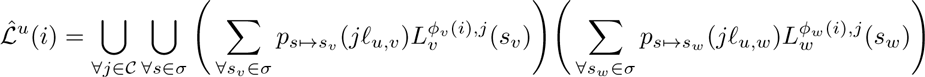

and the CLVs 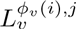 resp. 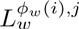 for all rates 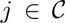 are retrieved from 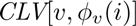 resp. 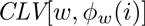.

**Figure 6.**
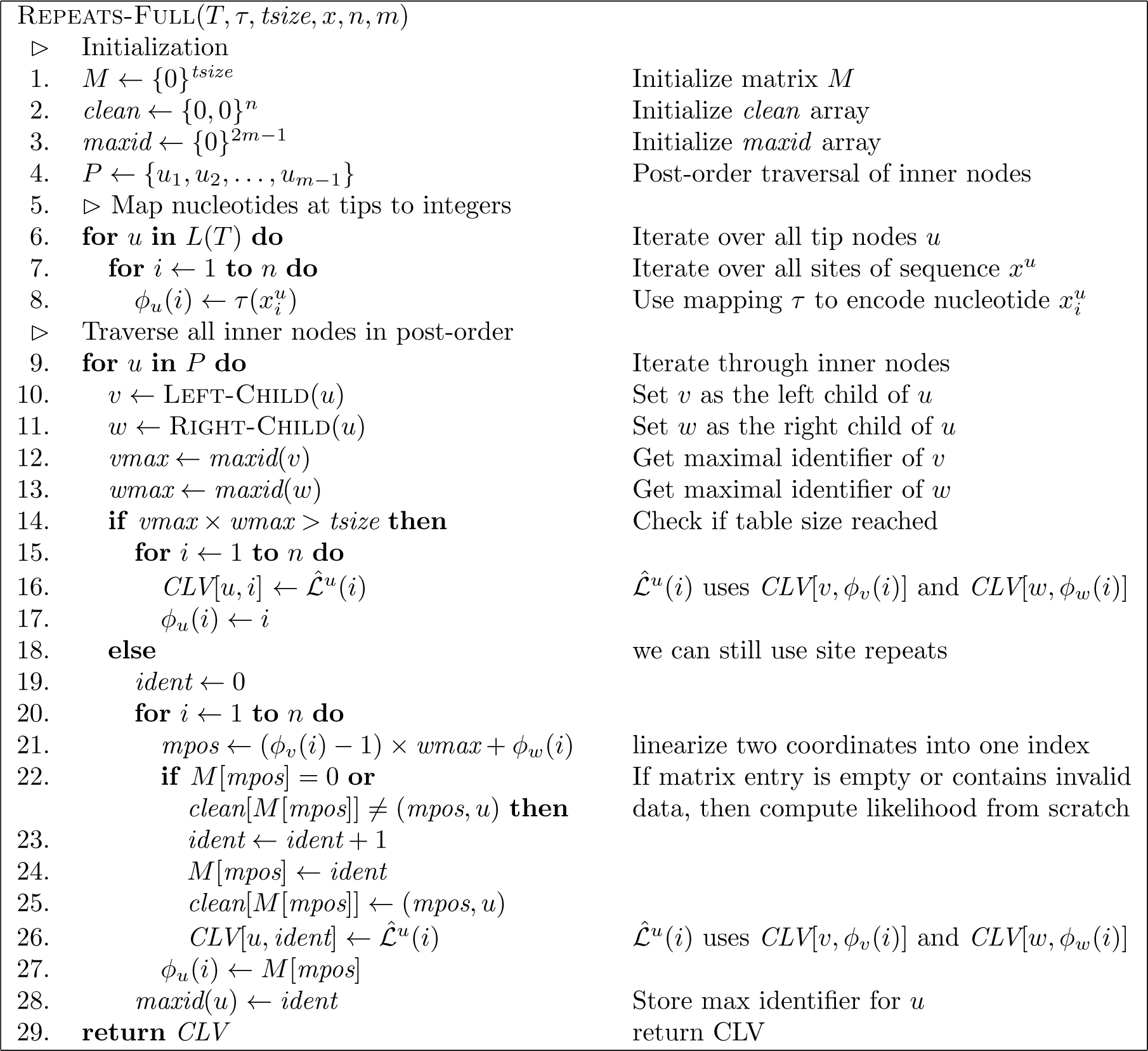
Full description for computing all CLVs of a tree *T* with the memory saving technique and site repeat detection. Input parameters are a tree *T* of *m* taxa, the sequences (of size *n*) for each of the *m* taxa (denoted 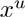 for the sequence at tip node *u*), a mapping *τ* for encoding the MSA data to integer values, and the size *tsize* of the matrix used for computing site repeats. The algorithm computes only the necessary CLVs required for evaluating the likelihood of tree *T*, and skips PLF calls on repeated sites.

**Observation 3 (Runtime)** *Algorithm* REPEATS-FULL *computes all site repeats, and the corresponding CLVs, in linear time with respect to the size of the alignment (number of sites times number of sequences), provided the allocated table M*.

This observation holds by inspection. For a description of the general, linear-time and *linear-space* algorithm that identifies all repeats in arbitrary *n*-ary trees and forests see Section 3a of [9]. Since that algorithm relies on several sorting steps using bucket sort — a linear-time sorting method not based on the comparison model —, in practice, it could be slower than the method we present here, which only queries a lookup table. To compute all repeats, our method requires quadratic space (with respect to the number of sites) in the worst case, which, however, is allocated only once and re-used for each node. When less memory is allocated, our method skips repeats identification for nodes where the product of unique site repeats for its two child nodes is larger than the entries of the allocated lookup table (i.e., low number of repeats). This is advantageous, since the overhead of repeat identification for such nodes could potentially cancel out the run-time savings.

## 3 Computational Results

We implemented a prototype of our algorithm in a new, low-level implementation of the PLL [7] (which we refer to as LLPLL), that does not make use of the highly optimized PLF of PLL, but allows for a straight-forward implementation of our algorithm. To demonstrate the applicability of our method under different settings, we created two implementations; one suitable for dynamically changing topologies (SRDT) and one suitable for fixed, constant tree topologies (SRCT). The first variant (SRDT) assumes no prior knowledge of the site repeats of a tree topology, and therefore, computes them before each PLF call. This variation is required for tree space exploration as site repeats change every time the tree topology is modified. The second variant (SRCT) computes site repeats only once as apart of an initialization step. Assuming a constant, fixed topology, the PLF reuses the precomputed information from the initialization step at each invocation, as site repeats remain unchanged. This variant is suitable for applications where no tree exploration is performed, as for example in divergence time estimation [11] and model selection [1], or during tree inference when parameters such as substitution rates or the α shape parameter of the gamma distribution are optimized.

To assess the performance of our method, we compared it with the sequential AVX-vectorized PLF implementation of the PLL which uses the same, highly optimized, PLF as RAxML. We selected PLL/RAxML because (i) it is our own code and hence we have a thorough understanding of it and (ii) it is currently among the fastest and most optimized PLF implementations available. This guarantees a fair comparison (i.e., determining the absolute speedup), and ensures that our method can truly be used in practice for speeding up state-of-the-art inference tools. We compare against two flavors of PLL; the plain version (we refer to it as PLL) and the memory saving SEV-based implementations of PLF (accessible using the `-U` switch in RAxML) which we refer to as PLL-SEV. The latter is faster and requires less memory than the former in the case of particularly gappy alignments [19]. In order to obtain an accurate speedup estimate of our method, we vectorized the LLPLL likelihood function using AVX instructions. However, since LLPLL is in an early development phase, the PLL is still faster by a factor of approximately 1.40 - 1.45 than LLPLL (without site repeats), as we show further. Despite this fact, we show that using our method, the LLPLL in its current state outperforms both PLL and PLL-SEV by up to a factor of 10. Our prototype implementation is available at http://www.exelixis-lab.org/web/software/site-repeats/.

**Data sets**. For performing the experiments we used a mixture of empirical and simulated DNA data sets which are summarized in Table 1. All data sets contain gaps and ambiguous DNA characters. Table 1 also reports the percentages of gaps and site repeats in the alignments. The amount of gaps is important, as it affects the performance of the PLL-SEV implementation. The percentages of site repeats are given for an arbitrary rooting of the parsimony trees calculated for these data sets using RAxML. While the data set with 2 000 taxa exhibits the lowest percentage of site repeats, it still has 86.95% repeats (which directly translate to identical conditional likelihood entries). We want to emphasize here, that we did not choose these data sets based on their repeat percentages. In fact, the fraction of site repeats for each data set was previously unknown to us. All data sets used for testing, are available on-line at (https://github.com/stamatak/test-Datasets).

**Table 1:** Summary of nucleotide data sets. For each data set, sites present the length of the provided MSA, and repeats denote the amount of sites over all nodes that are repeats of another site at the same node. The amount of repeats depends on the tree topology, the selected root and the MSA. The (unrooted) trees were obtained by running a maximum parsimony tree search for each of the data sets, and we randomly chose one node as the root to estimate the number of repeats.

|  |  |  |  |  |  |  |  |  |  |  |
| --- | --- | --- | --- | --- | --- | --- | --- | --- | --- | --- |
| **Data set/Sequences [-]** | 59 | 128 | 354 | 404 | 500 | 994 | 1 512 | 2 000 | 3 782 | 7 764 |
| **Sites [-]** | 6 951 | 29 198 | 460 | 13 158 | 1 398 | 5 533 | 1 577 | 1 251 | 1 371 | 851 |
| **Repeats [%]** | 92.04 | 91.78 | 94.65 | 96.49 | 89.43 | 94.63 | 90.09 | 86.95 | 94.18 | 87.62 |
| **Gaps [%]** | 44.24 | 32.48 | 14.71 | 78.92 | 2.25 | 71.39 | 3.02 | 12.65 | 2.70 | 20.60 |

**Experimental setup**. For assessing the performance of our method we conducted five types of experiments which cover the typical PLF use cases. First, we exhaustively assess the performance of full traversals for all possible rootings of the parsimony trees on two MSAs. Second, we assess the performance of full traversals on all ten MSAs for a limited number of random rootings. Third, we evaluate the performance for partial traversals, i.e. when not all CLVs need to be recomputed. Fourth, we assess the performance of our method on fixed tree topologies. In this setting, preprocessing of site repeats is done only once and not at each invocation of the PLF. Finally, using the three empirical multi-gene data sets in our collection, we determine the amount of memory required for maintaining the repeat tables when performing partitioned analysis. For the experiments we used a 4-core Intel i7-2600 multi-core system with 16 GB of RAM. To eradicate the potential impact of server-side events such as context-switching or performance peaks of running processes, we always executed several (usually 10 000) independent likelihood computations.

Considering that the calculation of the PLF takes up to 85%-98% of the total run-time of ML phylogenetic tree inferences [3], accelerating the performance of the PLF significantly impacts the overall execution time of ML analyses. Therefore, for the run-time comparisons we focus purely on the PLF evaluation. Branch lengths and model parameters are fixed, and remain unchanged as they do not impact the run-time of PLF.

Table 2 presents the memory savings due to site repeats together with the actual size of the lookup table for pre-processing all repeats. In the experiments, the size of the lookup table was bounded to 200 MB which corresponds to roughly 50 million entries (namely unsigned integer values). The actual memory for the lookup table was only allocated as needed. For most data sets, less than 200 MB of RAM was required. The notable exception is the data set containing 128 taxa, which requires 168.69 million entries (roughly 680 MB) to compute all site repeats. Since we bound the size of *M* to 200 MB, not all repeats were pre-processed when analyzing this particular data set.

**Table 2:** Summary of memory requirements for each method to evaluate the PLF at a random rooting of the parsimony tree. The table size entry specifies the size of the lookup table *M* of Algorithm REPEATS-FULLrequired to compute *all* possible repeats. It is presented in millions of entries (unsigned integers) and hence, its size in MB is four times as high as the presented numbers.

|  |  |  |  |  |  |  |  |  |  |  |
| --- | --- | --- | --- | --- | --- | --- | --- | --- | --- | --- |
| **Sequences** | 59 | 128 | 354 | 404 | 500 | 994 | 1 512 | 2 000 | 3 782 | 7 764 |
| **Sites** | 6951 | 29 198 | 460 | 13 158 | 1 398 | 5 533 | 1 577 | 1 251 | 1 371 | 851 |
| **Memory PLL [MB]** | 53 | 474 | 24.5 | 680 | 93 | 707 | 312 | 328 | 678 | 875 |
| **Memory PLL-SEV [MB]** | 46 | 403 | 21.5 | 326 | 93 | 256 | 308 | 297 | 674 | 819 |
| **Memory SRCT/SRDT [MB]** | 32 | 303 | 7.5 | 202 | 34 | 164 | 104 | 120 | 171 | 298 |
| **Table size** | 5.3 | 168.69 | 0.07 | 23.5 | 0.82 | 6.6 | 2.9 | 2.4 | 2.8 | 0.87 |

### 3.1 Exhaustive evaluation of all rootings

To get an initial estimate of the impact of distinct rootings on run-time, we used data sets 59 and 354 to evaluate the PLF at each *terminal* edge (an edge whose one end-point is a tip node) of their respective parsimony tree. This choice of rootings was selected because PLL requires that likelihood evaluations using full traversals of unrooted trees start at terminal edges. For each such rooting, we executed 10000 independent PLF computations using each of the four implementations: the LLPLL (without site repeats, SRDT (LLPLL with out method), PLL and PLL-SEV. For SRDT, we bounded the table size *M* to 200 MB, which, according to Table 2, is sufficient to find all repeats for these two data sets.

Table 3 summarizes the results of the experiments. PLL is, on average, 1.46 times faster than LLPLL on data set 59, and 1.39 times faster on data set 354. PLL-SEV has a slightly better runtime than PLL and is 1.58 times faster than LLPLL on data set 59 and 1.48 times faster on data set 354. The difference in speed between LLPLL and PLL/PLL-SEV can be explained by three factors. First, PLL is a highly optimized software for PLF calculations that was directly derived from RAxML, which in turn, has been developed and optimized for over 10 years, while LLPLL is in an early phase of development. Second, the standard optimization technique explained in the introduction, namely, the precomputation of conditional likelihoods for all combinations of two states with the subsequent querying from a lookup table (for tip-tip cases), is not implemented in LLPLL yet. This missing feature affects performance. Third, the LLPLL is designed to work with an arbitrary number of states which makes the (AVX vectorized) PLF slower than a dedicated 4-state PLF implementation. Despite this fact, SRDT is on average 3.59 times faster than PLL and 3.33 times faster than PLL-SEV for the “gappy” data set 59. Similarly, for data set 354, SRDT is on average 4.96 times faster than PLL and 4.65 times faster than PLL-SEV. Note that, the standard deviation for SRDT is higher than PLL (and PLL-SEV), which is expected given that the amount of repeats changes with different rootings.

**Table 3:**
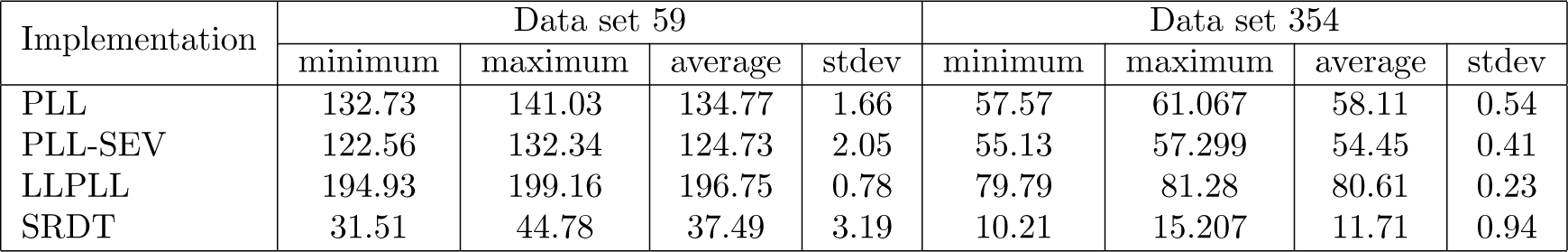
Summary of run-times for evaluating the PLF at each terminal edge of the parsimony trees (data sets 59 and 354). Presented are the minimum, maximum and average run-times over all rootings for each of the four implementations, and for each data set, along with the standard deviation of run-times among all rootings.

### 3.2 Evaluation of a sample of rootings

For the comprehensive comparison of full tree traversal times between SRDT and PLL, we use nucleotide data sets ranging from 59 to 7764 taxa (see Table 1). We measured the run-times for 10 different rootings chosen at random, and which are not necessarily the same for SRDT and the PLL implementations. However, this comparison is sufficient given the standard deviation among the run-times of different rootings computed for the exhaustive comparison of data sets 59 and 354. We again restricted the rootings to terminal branches, and for each of the 10 random rootings, we executed 10000 full tree traversals. For each of the three implementations (SRDT, PLL, and PLL-SEV) we computed the average run-time over all 10 rootings. Table 4 shows the speedup of SRDT compared to PLL and PLL-SEV. As we see, SRDT is consistently at least twice as fast as PLL. In fact, the lowest observed average speedup is 2.47. The maximal observed speedup was for the MSA with 404 taxa, where SRDT is 5.31 times faster than PLL. Table 1 shows that this particular data set has the highest relative number of repeats among all analyzed data sets, and signifies that the number of repeats positively affects the speedups over PLL. On the other hand, the largest decrease in speedups when comparing against PLL-SEV is observed for data sets 404 and 994 which comprise over 70% gaps. Note also, that run-times for PLL-SEV were higher than for PLL on data sets with a low amount of gaps, such as 1512, 3782, and 7764.

**Table 4:**
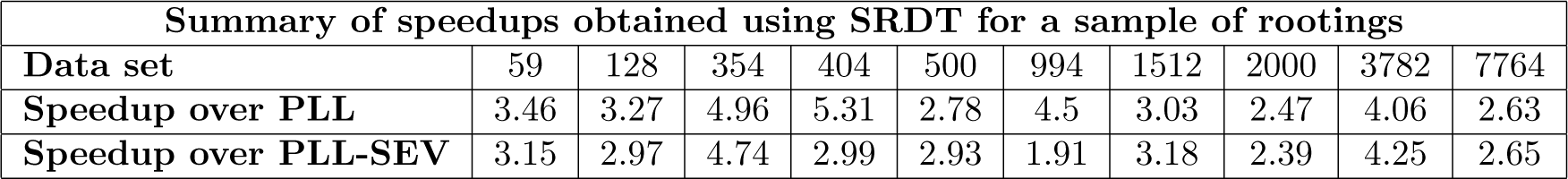
Speedups obtained when evaluating the PLF using SRDT over PLL and PLL-SEV for each of the ten data sets. SRDT is consistently faster than both methods. Speedups are computed as fractions of average run-times over 10 random rootings.

### 3.3 Partial traversal performance

In phylogenetic inference, it is not always necessary to perform full tree traversals to calculate the overall likelihood of a tree. In particular, when conducting BI or ML tree searches that deploy local topological updates using, for instance, nearest neighbor interchange (NNI) or subtree pruning and regrafting (SPR) moves, only the CLVs of a subset of tree nodes need to be recomputed. Depending on the topological update, those CLVs may be located at the inner part of the tree where the number of repeats is lower. We assessed the performance of our approach for this scenario by emulating partial CLV updates as follows. For each parsimony tree we choose two adjacent nodes (sharing an edge) at random, and place them in an empty list of nodes whose CLVs are to be updated. Next, with probability *p*, we choose whether to continue adding nodes to the list or stop (with probability 1 — *p*) the procedure. If we choose to add nodes, then we randomly pick one (unvisited) adjacent inner node to each of the (at most) two nodes selected in the previous step, and add them to the list. The procedure is repeated until we choose to stop or until no single inner node is available anymore for selection. This pattern of CLV updates emulates the topological moves described in [14] for BI. As mentioned before, in addition to the time spent in the PLF, other factors, such as branch length and model parameter optimization for ML, also contribute to the overall execution time. Here, we concentrate only on measuring the time for calculating the PLF. We used the method described above to simulate eleven partial CLV updates for each data set from Table 1. Each simulation consists of a path of nodes generated from a randomly chosen pair of adjacent nodes with probability *p* set to 0.95. We calculated each CLV along the path 10000 times and measured the total execution time for LLPLL and SRDT. Figure 7 presents the individual speedups for each data set and each simulation, plotted against the number of nodes updated for the particular simulation. We did not compare SRDT to PLL/PLL-SEV since a fair comparison requires that exactly the same partial updates are evaluated, which is difficult to achieve given the different internal structures of the two implementations. Therefore, the speedup for the partial updates is **not** the absolute speedup for different PLF implementations. Instead, our results show the relative speedup that can be achieved by incorporating our method into any PLF implementation.

**Figure 7.**
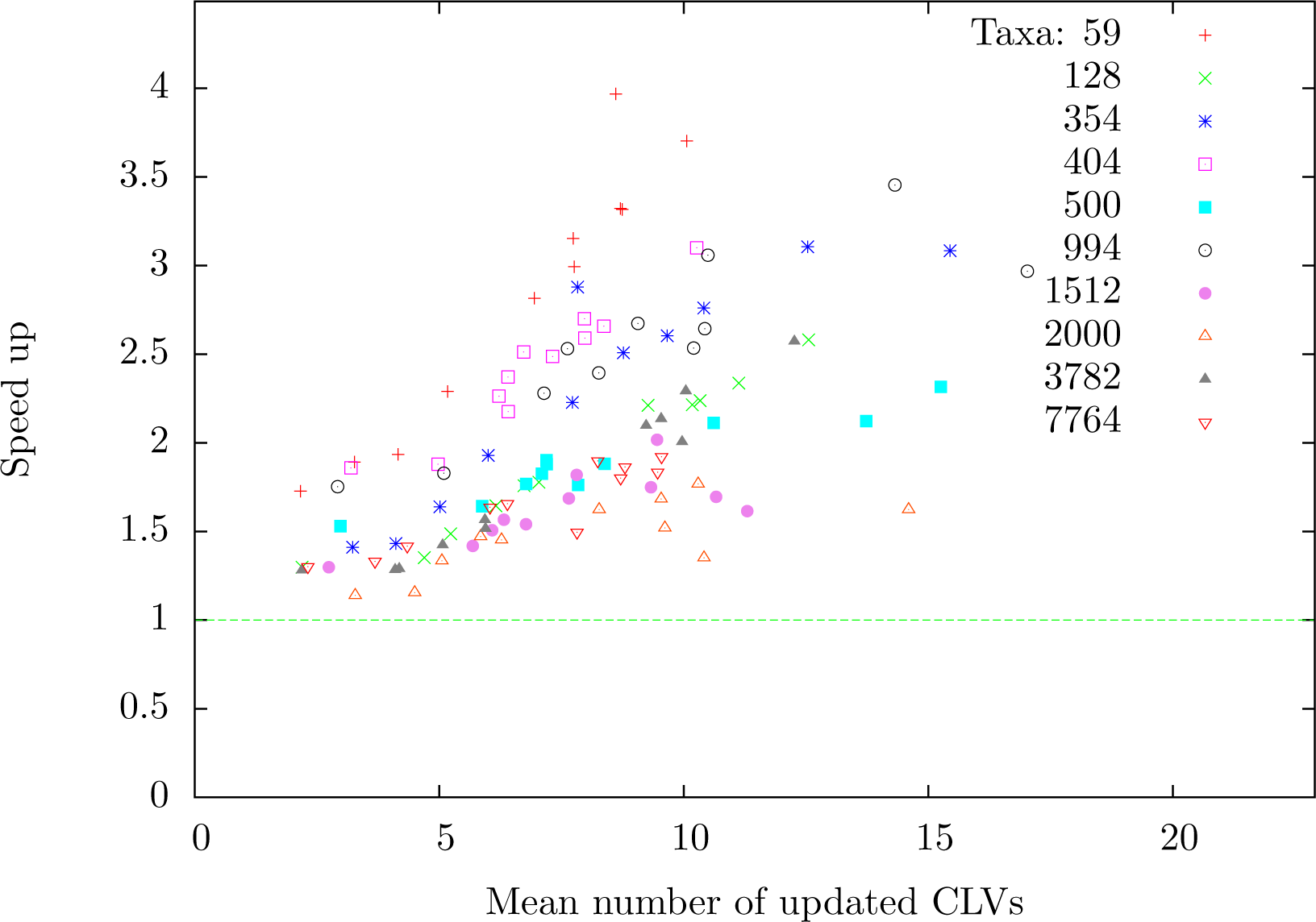
Run-time improvements of the SRDT method over the LLPLL method against the average number of updated CLVs. The colors distinguish the different data sets. Each data set is represented by eleven measurements for eleven different nodes.

### 3.4 Performance on fixed topologies

Many phylogenetic tools use a fixed tree topology on which the likelihood is repeatedly evaluated while other parameters, such as substitution rates and branch lengths, are modified. Divergence times estimation [11] and model selection [1] are typical representations for this setting. With fixed topologies, repeats can be precomputed once in an initialization step, and then reused for subsequent PLF invocations. We tested our method under this setting, by evaluating the likelihood 10000 times, rooted at 10 randomly selected terminal edges of the parsimony trees of each of the 10 data sets. We averaged the run-time over the 10 rootings and report the speedups of SRCT over PLL and PLL-SEV in Table 5. We computed the speedup for each data set as the ratio of the average run-time of PLL resp. PLL-SEV from Section 3.2 (evaluation of a sample of rootings) divided by the average run-time of SRCT. Data set 994 yields the lowest speed up over PLL-SEV, which is due to the high number of gaps in the data set, followed by data set 2000 which has the lowest number of repeats. On the other hand, the highest speedup of SRCT over PLL-SEV was for data set 354 which has a high number of repeats combined with a low amount of gaps. As expected, the highest speedup of SRCT over PLL is observed for data set 404, since it has the highest amount of repeats among all data sets.

**Table 5:** Speedups obtained using the SRCT method which considers a fixed topology over the PLL and PLL-SEV.

| Summary of speedups obtained using SRCT when considering fixed topologies |  |  |  |  |  |  |  |  |  |  |
| --- | --- | --- | --- | --- | --- | --- | --- | --- | --- | --- |
| Sequences | 59 | 128 | 354 | 404 | 500 | 994 | 1512 | 2000 | 3782 | 7764 |
| Sites | 6951 | 29198 | 460 | 13158 | 1398 | 5533 | 1577 | 1251 | 1371 | 851 |
| Speedup over PLL | 5.71 | 4.64 | 8.59 | 9.96 | 4.29 | 8.16 | 4.66 | 3.62 | 6.86 | 3.91 |
| Speedup over PLL-SEV | 5.22 | 4.23 | 8.22 | 5.44 | 4.52 | 3.30 | 4.88 | 3.48 | 7.18 | 3.93 |

### 3.5 Partitioned analyses

We performed partitioned analysis on the three empirical multi-gene data sets (59, 128 and 404) in our collection to compare the accumulated memory requirement of per-partition repeat tables against single repeat tables from unpartitioned analyses. This experiment is of high practical relevance because typical phylogenetic analyses are partitioned nowadays. Table 6 provides a summary of the per-gene partitions of each data set, the required per-partition table size for computing all repeats, and the per-partition speedups over PLL-SEV. For measuring the speedups, we used the same parsimony trees from the previous experiments, and computed an average run-time over 10 random terminal edge rootings. As in the rest of the experiments, the rootings may not be the same between PLL-SEV and LLPLL. Similarly to Table 2, per-partition table sizes are presented in millions of entries (unsigned integers). One important result is that the accumulated table size for each partitioned data set is considerably smaller than the table size required for processing single-partitioned data sets (see Table 2). Compared to their single-partition counterparts, dataset 59, 128 and 404, require 10, 13, and 6 times less memory for storing the per-partition tables. Not surprisingly, this is due to the way we index elements in the table (see line 21 of Figure 6). This kind of indexing may cause the table to grow quadratically to the maximum number of unique repeats at a node, which in turn, increases with the size of the alignment.

**Table 6:** Summary of per-partition table sizes for data sets 59, 128, and 404. All table sizes are presented in millions of entries (unsigned integers) and indicate the size of the lookup table *M* required to compute all possible repeats. When performing partitioned analysis, data set 59 requires a total of 0.521 million entries, data set 128 requires 12.88 and data set 404 requires 3.74. We also present the speedups over PLL-SEV for evaluating the PLF for each partition.

| Dataset 59 |  |  |  |  |  |  |  |
| --- | --- | --- | --- | --- | --- | --- | --- |
| Partition | Sites | Speedup | Table size | Partition | Sites | Speedup | Table size |
| 5.8S | 160 | 5.05 | 0.001 | ndhf1st | 2 183 | 3.53 | 0.24 |
| cprs | 364 | 2.08 | 0.053 | phyb3rd | 1 182 | 3.14 | 0.08 |
| gbss13rd | 774 | 3.56 | 0.016 | rbcl1st | 1 344 | 4.93 | 0.033 |
| its2 | 264 | 1.67 | 0.06 | rpoc23rd | 680 | 2.59 | 0.038 |
| Dataset 128 |  |  |  |  |  |  |  |
| Partition | Sites | Speedup | Table size | Partition | Sites | Speedup | Table size |
| 12S_rRNA | 860 | 3.27 | 0.28 | Cytb | 1 149 | 2.70 | 0.74 |
| 16S_rRNA | 1 275 | 3.40 | 0.36 | EDG1 | 978 | 4.92 | 0.085 |
| ADORA3 | 330 | 3.68 | 0.025 | IRBP | 1 233 | 3.74 | 0.45 |
| ADRB2 | 833 | 4.78 | 0.08 | ND1 | 963 | 2.80 | 0.56 |
| APOB | 1 335 | 3.67 | 0.47 | ND2 | 1 050 | 2.45 | 0.85 |
| APP | 717 | 3.88 | 0.09 | ND3 | 349 | 2.27 | 0.1 |
| ATP6 | 681 | 2.41 | 0.24 | ND4L | 297 | 2.27 | 0.057 |
| ATP7A | 684 | 3.94 | 0.095 | ND4 | 1 378 | 2.52 | 0.99 |
| ATP8 | 213 | 1.96 | 0.048 | ND5 | 1 836 | 2.41 | 2.69 |
| BDNF | 582 | 4.88 | 0.024 | ND6 | 555 | 2.36 | 0.23 |
| BMI | 340 | 4.82 | 0.0086 | PLCB4 | 373 | 2.82 | 0.067 |
| BRCA1 | 2 934 | 3.27 | 2.38 | PNOC | 333 | 3.64 | 0.022 |
| CNR1 | 993 | 5.58 | 0.069 | RAG1 | 975 | 4.43 | 0.088 |
| COX1 | 1 557 | 3.27 | 0.77 | RAG2 | 444 | 4.54 | 0.024 |
| COX2 | 708 | 2.69 | 0.22 | tRNAs | 1 341 | 3.20 | 0.46 |
| COX3 | 804 | 2.76 | 0.24 | TYR | 426 | 3.80 | 0.028 |
| CREM | 468 | 3.42 | 0.04 | ZFX | 204 | 4.96 | 0.0022 |
| Dataset 404 |  |  |  |  |  |  |  |
| Partition | Sites | Speedup | Table size | Partition | Sites | Speedup | Table size |
| ndhF | 2 161 | 3.62 | 1.17 | rbcl | 1 437 | 4.65 | 0.28 |
| rpoc2 | 802 | 2.39 | 0.24 | phyB | 1 218 | 3.25 | 0.28 |
| tRNA | 1 646 | 2.39 | 1.15 | psbA | 1 020 | 3.44 | 0.002 |
| rps4 | 588 | 5.20 | 0.0027 | rps16 | 1 025 | 2.62 | 0.032 |
| adh1 | 852 | 3.01 | 0.03 | matK | 1 581 | 4.10 | 0.22 |
| its1,5.8S,its2 | 828 | 2.22 | 0.33 | - | - | - | - |

## 4 Conclusion

The PLF is among the computationally most time-consuming functions in evolutionary biology, and typically constitutes the largest portion of total run-time time in analyses involving PLF calculations. Especially in the era of genomics, where datasets comprise thousands or even millions of alignment sites [13, 15], accelerating the PLF can save weeks or even months of CPU time.

We introduce a novel method for quickly identifying all repeating site patterns, and consequently, minimizing the number of operations required for evaluating the PLF. It is based on our linear-time and linear-space algorithm for identifying repeating subtrees in general labeled, unordered, *n*-ary trees. Our new method is optimized to work with phylogenetic trees (rooted or unrooted binary trees) and discards many of the hidden constants behind the complexity of the original method for *n*-ary trees.

To measure the speed of the PLF during tree searches or model parameter optimizations, we compared a prototype implementation of our method against PLL - a library derived from the RAxML code — which uses one of the fastest and most highly tuned implementations of the PLF. Using empirical and simulated data, we measured the speedup under different, realistic settings. For fixed and dynamically changing tree topologies we observe an up to 10-fold speedup. For partial CLV updates, i.e., when only a small number of CLVs is recomputed due to a topological rearrangement of the tree, we still observe a 2- to 5-fold speedup. Moreover, our method, *including* the book-keeping information for site repeats requires significantly less memory than PLL, sometimes up to 78% less. Using empirical data, we show that the book-keeping storage requirements for partitioned analyses are significantly smaller than for unpartitioned analyses (up to 13 times less). This result is particularly important for optimizing large phylogenomic analyses. Our method is simple, and can be incorporated into any implementation of the PLF. Moreover, the table used for book-keeping information is flexible, and could be dynamically adjusted, for example in an “auto-tuning” step, to determine the table size — and consequently the amount of identified repeats — that minimizes PLF runtime. Tree nodes with a number of unique repeats that is close to the number of sites, i.e., with a small amount of repeated sites, have higher memory requirements for book-keeping. When the allocated table size is exceeded our method will omit repeat identification for all subsequent nodes, as the amount of repeats decreases towards the root of the tree. This allows our method to omit repeat computations at nodes for which calculating them is disadvantageous.

